## Supplemental Information for "Using experimental results of protein design to guide biomolecular energy-function development"

**Table of contents**

Supplemental Tables - pg. 2-4
Supplemental Figures - pg. 5-16
Supplemental Text - pg. 17-18

### Supplemental Tables

| study | type of design | name of design | PDB ID of crystal structure |
|---|---|---|---|
| Chen et al., 2019, Nature [1] | helical heterodimer | DHD127 | 6DLM |
| | | DHD131 | 6DKM |
| Hicks et al., 2022, PNAS [2] | helical homodimer | D_3_212 | 7RMX |
| | | D_3_633 | 7RKC |
| Brunette et al., 2015, Nature [3] | helical repeat protein | DHR4 | 5CWB |
| | | DHR76 | 5CWO |
| Xu et al., 2020, Nature [4] | helical transmembrane pore | WSHC6 | 6TMS |
| Boyken et al., 2016, Science [5] | helical homotrimer | 5L6HC3_1 | 5IZS |
| Lajoie et al., 2020, Science [6] | helical bundle | 6AYYA | 7JH5 |
| Huddy et al., 2024, Nature [7] | helical bundle | THR1 | 8G9J |
| | | THR2 | 8G9K |
| Wu et al., 2023, Nature [8] | peptide binder | RPB_PLP3_R6–PLPx6 | 7UE2 |
| Sahtoe et al., 2022, Science [9] | $\alpha/\beta$ heterodimer | LHD29 | 6WMK |
| Basanta et al., 2020, PNAS [10] | NTF2 ($\alpha/\beta$ monomer) | BBM2nHm0589 | 6W3D |
| | | CAV1 | 6W3W |
| | | MC2_7 | 6W40 |
| Dou et al, 2018, Nature [11] | $\beta$ barrel | BB1 | 6D0T |

**S1 Table. List of crystal structures of designs analyzed in this study.** See https://github.com/Haddox/design_guided_optE/tree/main/designs_and_xtals for PDBs for each design model and its corresponding crystal structure, named according to this table. Most designs in the table have helical topologies, though a few have mixed $\alpha/\beta$ topologies, and one is a $\beta$ barrel.

| parameter | | beta_nov16 | beta_jan25 |
|---|---|---|---|
| | fa_rep | 0.5500 | 0.6191 |
| LJ radius ( Å ) | CH0 | 2.012 | 2.040 |
| | CH1 | 2.012 | 2.010 |
| | CH2 | 2.012 | 2.010 |
| | CH3 | 2.012 | 2.060 |
| | aroC | 2.016 | 2.020 |
| | Hapo | 1.421 | 1.450 |
| | Haro | 1.375 | 1.410 |
| LJ well depth (kcal/mol) | CH0 | 0.063 | 0.075 |
| | CH1 | 0.063 | 0.062 |
| | CH2 | 0.063 | 0.064 |
| | CH3 | 0.063 | 0.064 |
| | aroC | 0.069 | 0.070 |
| | Hapo | 0.022 | 0.021 |
| | Haro | 0.016 | 0.021 |

**S2 Table. Differences in parameter values between beta_nov16 and beta_jan25.**

| **Training Parameter** | **Value** | **Description** |
|---|---|---|
| Number of trees | 1000 | The number of independent decision trees trained per model. |
| Splitting criterion | Squared error | The quantity used to determine the optimal value at which to split a feature coordinate into two regions. The squared difference between the mean in each terminal node and its members (i.e., the variance). |
| Minimum samples per leaf | 20 | The minimum number of samples required in a leaf node. A split point at any depth was considered if it left at least 20 training samples in each of the left and right branches. |
| Maximum features per tree | 0.5 | The fraction of the total number of features considered when looking for the best split. At each split, a random selection of 50% of the features was considered to split. |
| Max depth | None | A maximum depth to which trees were allowed to be built. No restriction was placed on the depth of trees, instead depth was controlled implicitly using the *minimum samples per leaf* parameter. |
| Minimum impurity decrease | 0.0 | No threshold was placed on the minimum amount the squared error needed to be reduced to allow a split to occur. |
| Maximum number of leaf nodes | None | No explicit restriction was placed on the number of allowable leaf nodes. |
| Bootstrap | False | The whole training set was used to build each tree. |
| Minimum weight fraction per leaf | 0.0 | Samples were not weighted, so no minimum fraction of the total weight represented in leaf nodes was used to constrain leaves. |

**S3 Table**. Random forest regression model training parameters used for all topology-specific models. See [13] for further details on the random forest implementation used. See https://github.com/Haddox/design_guided_optE/tree/main/id_outlier_designs and https://zenodo.org/records/17330114 for data and code used to train models and identify outlier designs.

### Supplemental Figures

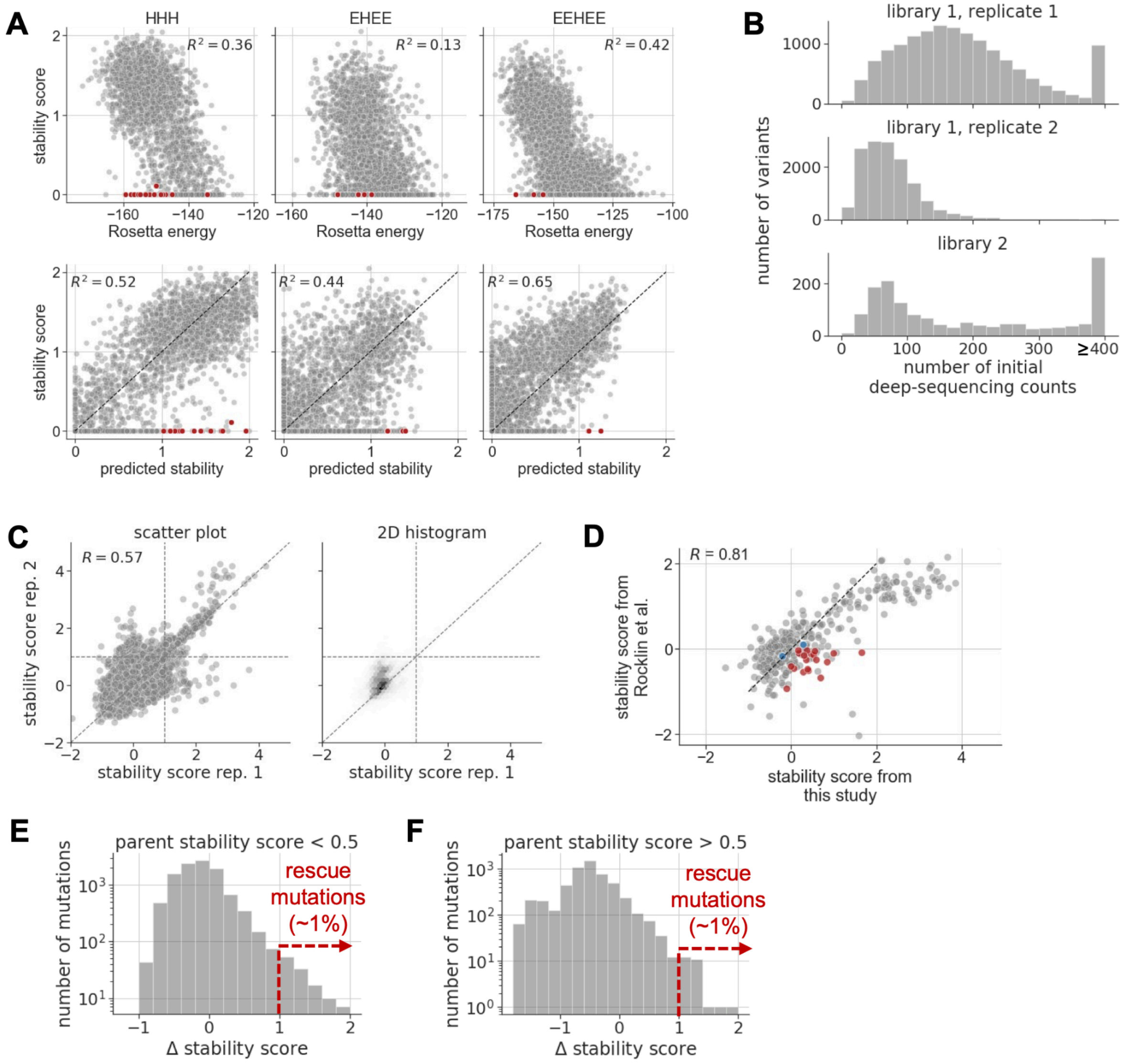


**S1 Fig. Additional results from deep mutational scanning of outlier designs. A)** The top row of plots are similar to Fig 2A, but show data for all three miniprotein topologies from Rocklin et al. that we analyzed in this study (HHH, EHEE, and EEHEE; letters indicate the order of secondary-structure elements in a topology, with H=helix and E=strand). The bottom row of plots are the same as the top row, but show stabilities predicted by ML models instead of Rosetta (see *Methods*). Red dots show all 21 outlier designs selected for DMS, which span all three topologies. **B)** Histograms show the number of deep-sequencing counts of all single amino-acid variants in starting DMS libraries after transforming them into yeast cells. As described in the *Methods*, the 21 DMS libraries were ordered and tested as part of two larger libraries. The first of these libraries was tested in duplicate. Each plot shows data for a single replicate of a given library. Distributions are capped at 400 counts. **C)** Correlation of stability scores between the two biological replicates of library 1, which were generated from independent transformations of unselected plasmid libraries into yeast. In the scatter plot (left), each dot corresponds with a unique variant in the library. The 2D histogram (right) shows that most variants had low stability scores near zero in each replicate. **D)** Correlation of stability scores between this study and Rocklin et al. for all outlier designs, as well as 338 “ladder” miniproteins from Rocklin et al. (see *Methods*). Red dots show outlier designs from library 1, blue dots show outlier designs from library 2, and gray dots show ladder miniproteins. *R* is the Pearson correlation coefficient. The higher dynamic range of values in this study comes from experimental modifications that made the yeast surface-display scaffolding proteins less sensitive to proteolysis (see *Methods*). **E)** The same as Fig 2C, but with the y-axis scaled in log space for increased visibility of the tails of the distribution. **F)** The same as panel E, but now showing mutational effects pooled across the 7 outlier designs with a stability score >0.5, as measured in this study.

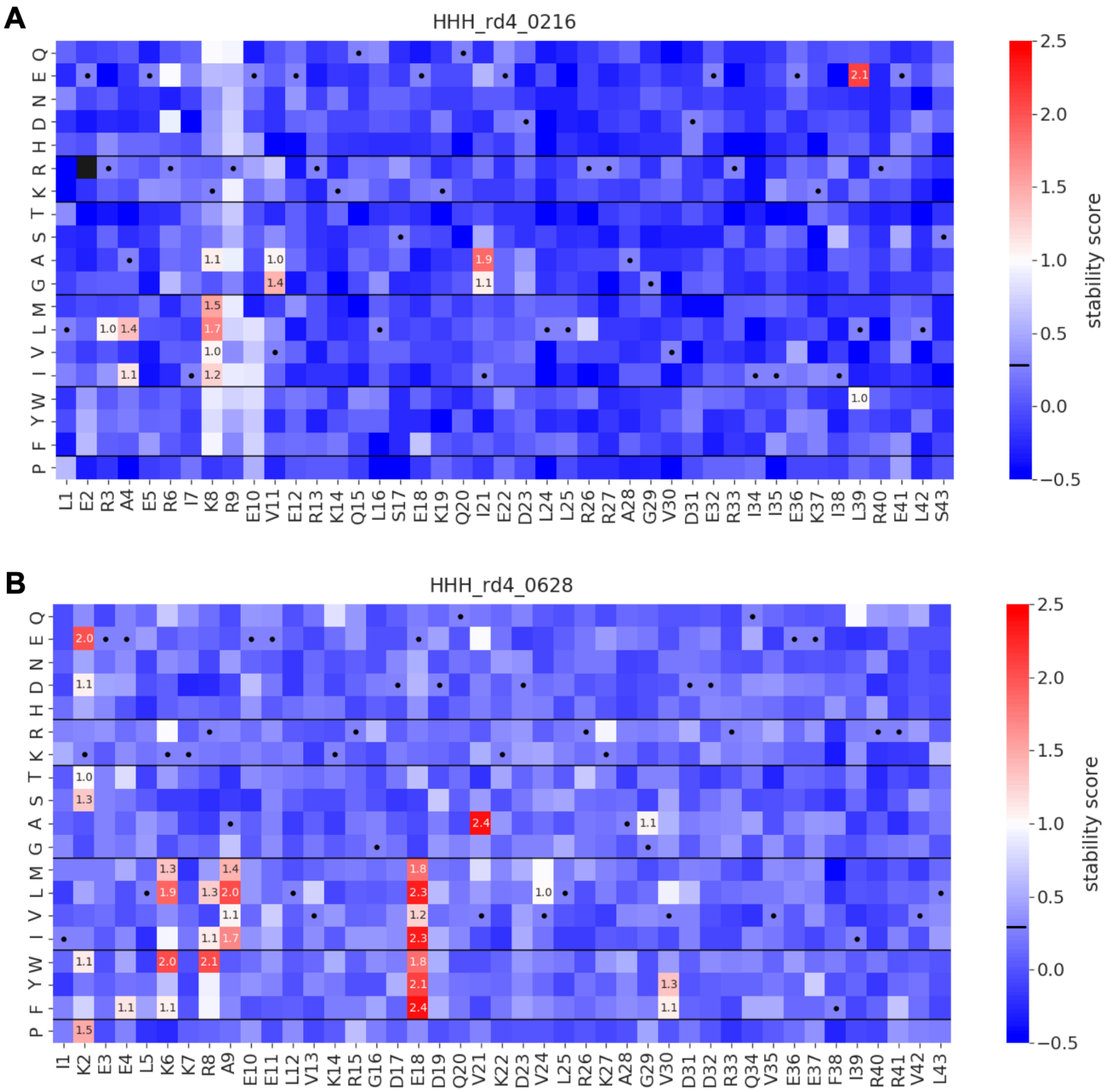


**S2 Fig. Stability scores of all single amino-acid variants from deep mutational scanning of two of the unstable outlier designs. A)** Data for HHH_rd4_0216 (labeled with an orange triangle in Fig 2G). **B)** Data for HHH_rd4_0628 (labeled with a purple circle in Fig 2G). Each box shows the stability score of a variant with a given amino-acid mutation (y-axis) at a given site (x-axis). Boxes with dots correspond to the unmutated design and show its stability score (the horizontal black line on the color bar also shows the unmutated design's score). Numbers show stability scores for variants with scores of at least 1.0. We were unable to measure stability scores for a small number of variants (black boxes). Heatmaps of all 21 outlier designs are available at https://github.com/Haddox/design_guided_optE/tree/main/rescue_dms/heatmaps.

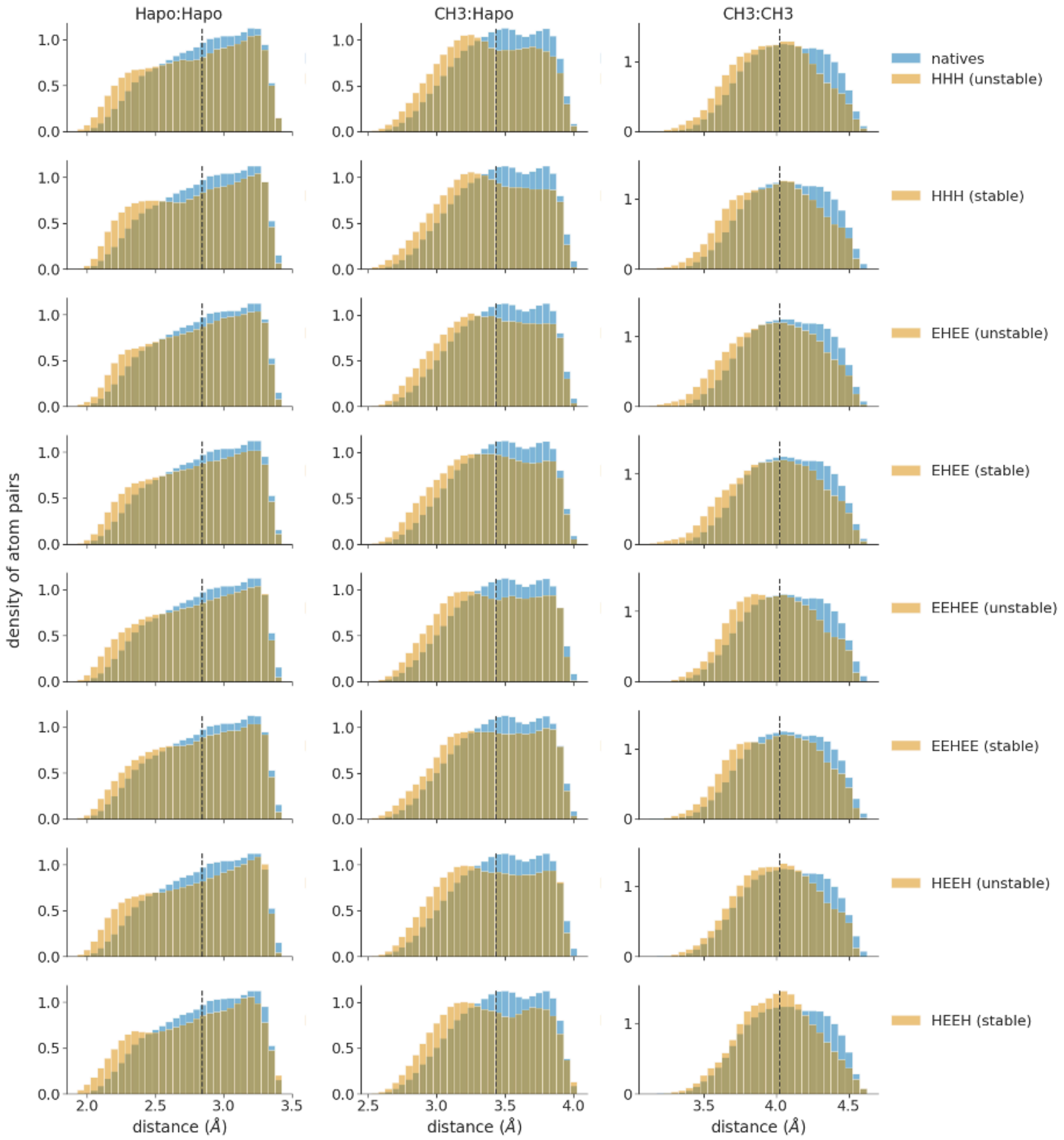


**S3 Fig. Levels of clashing in miniprotein designs exceed levels seen in high-resolution crystal structures of native proteins.** Each plot shows smoothed distributions of interatomic distances for a given atom pair (columns) within a given set of proteins (rows). Blue distributions show distances observed in 78 high-resolution crystal structures, showing data for all instances of a given atom pair within a distance cutoff of 0.5 Å greater than the sum of the radii of these atoms. Orange distributions show distances observed in a given set of designs from Rocklin et al. (from design rounds 3 and 4), with rows of plots separating designs by topology (HHH, EHEE, EEHEE, HEEH) and whether the designs were stable (stability score > 1.0) or unstable (stability score < 1.0). Vertical dashed lines show the sum of the van der Waals radii of the atom pair. Interatomic distances to the left of this line correspond to “clashes”. Columns show data for different atom pairs from hydrophobic sidechains, including pairs of hydrogens (Hapo:Hapo), hydrogens and methyl carbons (CH3:Hapo), or methyl carbons (CH3:CH3). In each plot, designs show higher levels of clashing than the crystal structures.

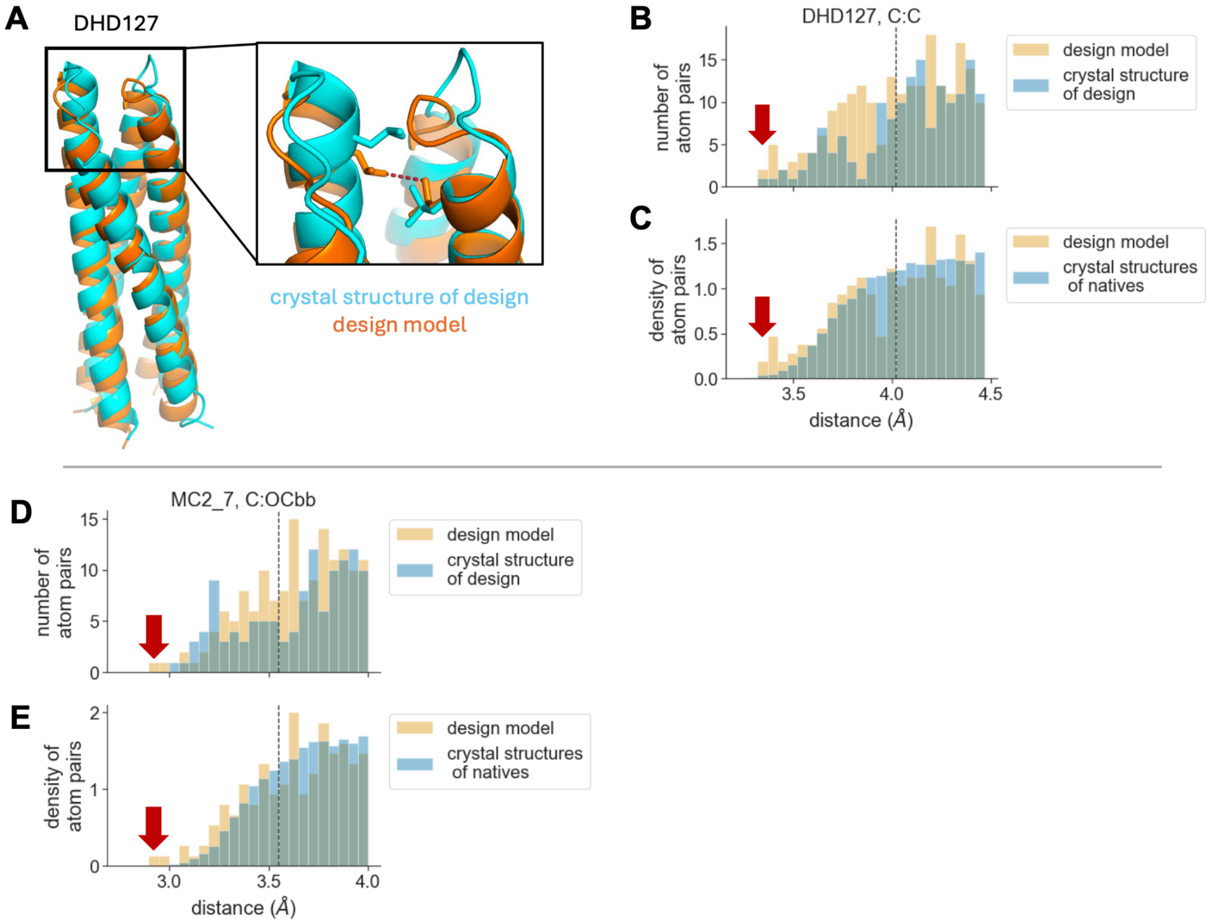


**S4 Fig. Clashing in design models compared to corresponding crystal structures. A)** An example of a large clash between carbon atoms from nonpolar sidechains (see dashed red line) in a designed heterodimer (DHD127; PDB ID 6DLM) [1]. The clash is present in the design model (orange), but absent in the crystal structure (blue) due to a conformational rearrangement in the region of the clash, where one of the clashing sidechains shifts up to occupy space previously occupied by a loop. The clash is more extreme than most clashes in the design's crystal structure and in a set of 54 crystal structures of native proteins, as shown in the next two panels. **B)** Histograms show the distribution of interatomic distances between all pairs of carbon atoms from nonpolar sidechains (C:C) in the DHD127 crystal structure (blue distribution) or design model (orange distribution) within a distance cutoff of 0.5 Å greater than the sum of the radii of these atoms (the dashed line shows this sum). The red arrow shows the distance of the clashing atom pair highlighted in panel A. **C)** Same as panel B, but the blue distribution shows interatomic distances from a set of 54 high-resolution crystal structures of native proteins. The two distributions are plotted as normalized densities to make them comparable. **D and E)** These three panels are similar to panels B and C, but show data for a designed NTF2 from Fig 3C, and examine clashing between pairs of carbon atoms from nonpolar sidechains and backbone oxygen atoms (C:OCbb). The red arrow shows the distance of the clashing atom pair highlighted in Fig 3C. See https://github.com/Haddox/design_guided_optE/tree/main/designs_and_xtals/compute_interatomic_distances/distance_distribution_plots for similar plots for all design-crystal pairs.

**A**

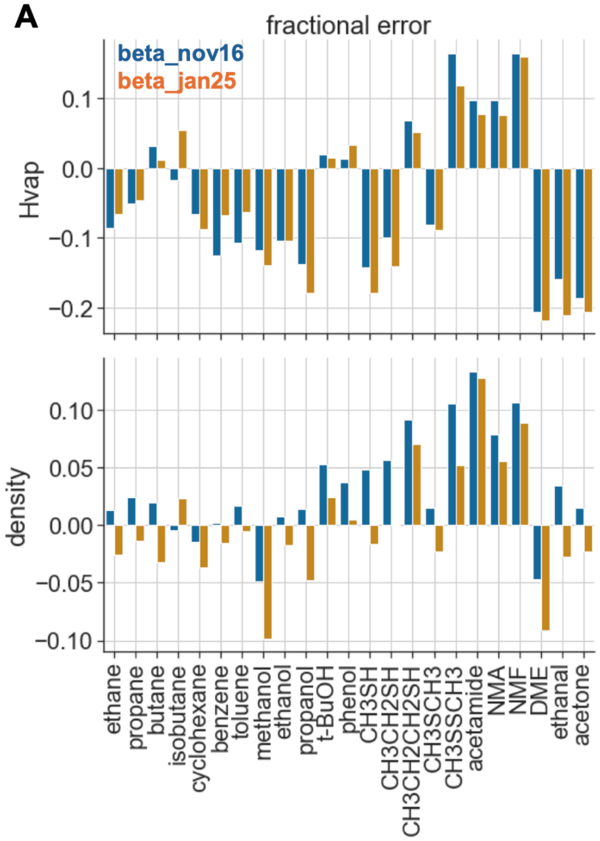


**B**

| energy function | average fractional error | |
|---|---|---|
| | **Hvap** | **density** |
| beta_nov16 | 0.10 | 0.04 |
| beta_jan25 | 0.10 | 0.04 |

**S5 Fig. Energy-function performance on the benchmark for predicting thermodynamic properties of small molecules.** For a panel of small molecules, the benchmark uses a liquid-simulation framework to predict each molecule's heat of vaporization and density, given an input energy function. We performed this benchmark as described in Park et al. [12]. **A)** Fractional error in predicted values for heat of vaporization (Hvap) and density. Each bar shows the fractional error for a given energy function (hue) on a specific small molecule (x-axis). Values are averaged over predictions from four replicate simulations for beta_nov16 or three replicate simulations for beta_jan25. (NMA = N-methylacetamide, NMF = N-methylformamide, DME = dimethyl ether). **B)** The average fractional error of each energy function in predicting the properties from panel A.

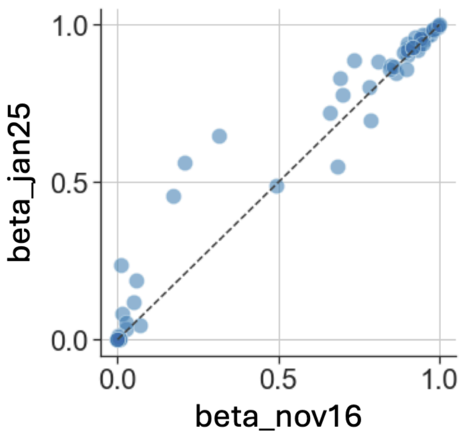


**S6 Fig. Energy function performance in the interface-prediction benchmark.** Each dot corresponds to one of the 59 crystal structures withheld from training. For each of these crystal structures, we generated a large number of decoys with non-native interfaces, relaxed the decoys with a given energy function, and then computed the Boltzmann-weighted probabilities of observing near-native structures (see *Methods*), with higher probabilities indicating better performance. The x and y axes report probabilities obtained by relaxing the structures with either beta_nov16 or beta_jan25, respectively. Table 1 reports the average probability for a given energy function. The higher average for beta_jan25 comes from modest increases in probabilities for several structures (dots above the diagonal line). There are only a few structures where the probability is lower for beta_jan25 (dots below the diagonal line).

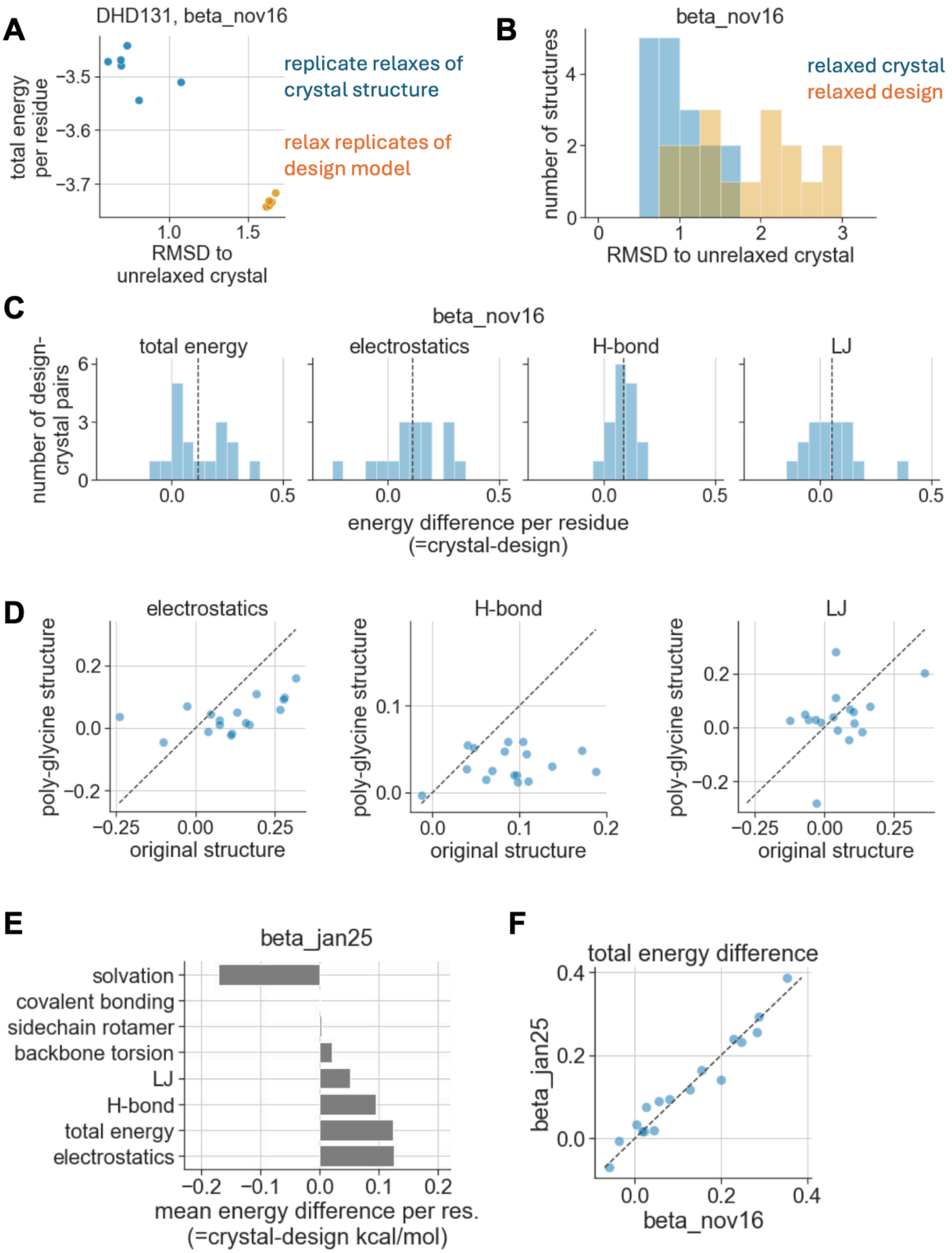


**S7 Fig. Energy differences between design models and corresponding crystal structures.** **A)** We separately relaxed each design model and each crystal structure, performing six independent replicates of the relax protocol. This panel shows the results for one design-crystal pair (DHD131) relaxed with the beta_nov16 energy function. Each dot corresponds the output structure from a single replicate of relaxing the crystal structure (blue dots) or the design model (orange dots). The y-axis shows each structure's total energy per residue, while the x-axis shows each structure's $C_\alpha$ RMSD to the unrelaxed crystal structure. See https://github.com/Haddox/design_guided_optE/tree/main/designs_and_xtals/relax_and_scoring_protocols/energy_landscape_plots for similar plots for all design-crystal pairs. For a given pair, we selected the lowest-energy replicate from each color category for the comparisons in Fig 5A and for the other panels in this figure. **B)** The $C_\alpha$ RMSD of relaxed design models (orange) and crystal structures (blue) to the unrelaxed crystal structure of the corresponding design. This plot shows data across all design-crystal pairs. **C)** Distributions of energy differences of design-crystal pairs. Vertical dashed lines show the mean of each distribution. Fig 5A plots the averages of these distributions. **D)** For each relaxed design model and each relaxed crystal structure, we created a version of the structure where all side-chain atoms were replaced with a single hydrogen atom (converting the sequence to poly-glycine while keeping all backbone atoms fixed in space). This panel shows energy differences computed using the unmodified structures with all side-chain atoms intact (full pose) compared with the poly-glycine structures (poly-Gly pose), with one dot for each design-crystal pair. Many of the energy differences are closer to zero for the poly-glycine structures. **E)** The same as Fig 5A, but for design-crystal pairs relaxed with the beta_jan25 energy function. **F)** The total energy differences of design-crystal pairs are highly correlated between energy functions.

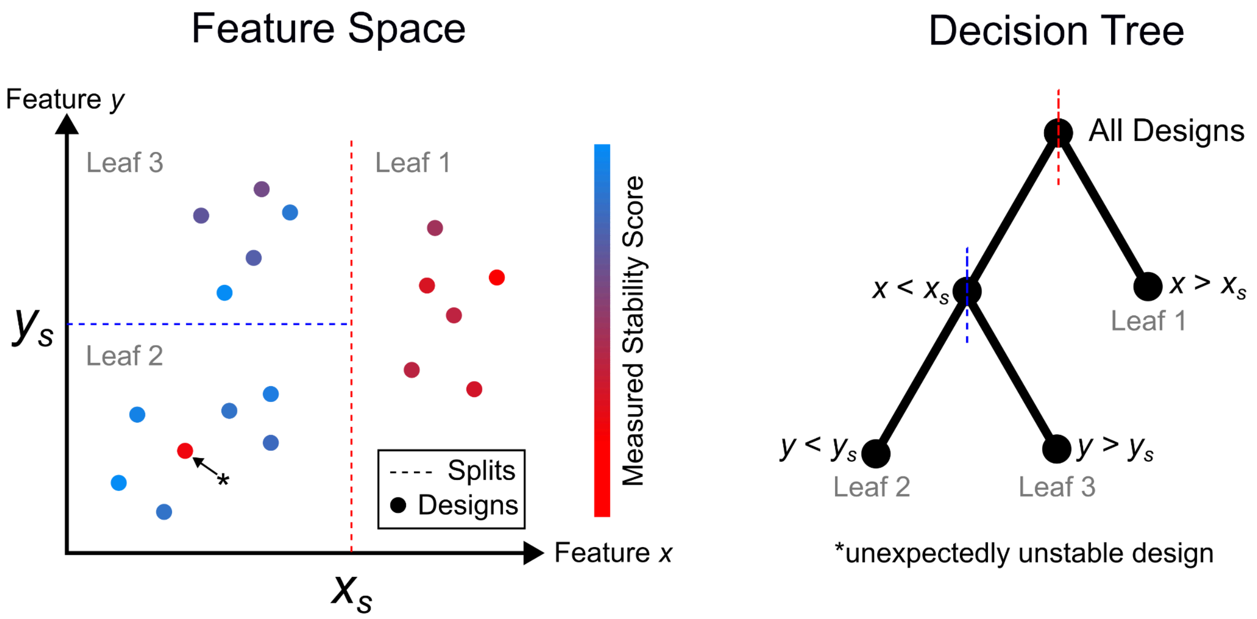


**S8 Fig. Random forest model and outlier design illustration.** Regions in a hypothetical 2-dimensional biophysical feature space determined by two splits resulting in a depth-2 decision tree with three leaves. Designs (points in feature space) are colored by cartoon experimentally measured stability scores. The indicated design with a low measured stability score (red point) in Leaf 2 will exhibit a large (negative) difference between its measure stability score and its model-predicted score since its leaf-neighbors show consistently high stability scores. Thus it may be identified as an unexpectedly unstable outlier design in this feature space.

### Supplemental Text

#### Random forest model training and data-driven identification of outlier designs

We examined miniprotein designs from Rocklin et al. [14], specifically focusing on designs from the HHH, EHEE, and EEHEE topologies. We filtered out designs with $EC_{50}$ 95% confidence intervals that spanned more than 2 units, helping to eliminate designs with substantial uncertainty in experimentally measured stabilities. A total of 10,232 designs met these criteria (2,304 for HHH, 2,965 for EHEE, and 4,963 for EEHEE).

We fit three random forest regression models, one for each miniprotein topology (HHH, EHEE, or EEHEE), using scikit-learn [13] (S3 Table). We trained each model to predict the experimentally measured stability scores of designs from a given topology using ~1,500 biophysical features computed for each design (the number of features varied slightly between topologies; see https://github.com/Haddox/design_guided_optE/tree/main/id_outlier_designs and https://zenodo.org/records/17330114 for the full set of features used for each topology). The features are an expanded set of the ones used to train models in Rocklin et al., and include the Rosetta energy of each design. Prior to training, each feature was standardized to have a mean 0 and standard deviation 1 for designs from a given topology. Models were trained to predict experimentally measured stability scores that were generated using the unfolded state model described in Singer et al. [15]; these stability scores are highly correlated with the ones originally reported in Rocklin et al.

To derive a predicted stability score for each design, designs with the same miniprotein topology were divided into 50 disjoint training/predicting splits so that approximately 98% of designs were used to train a random forest model that was subsequently used to predict the stability scores of the remaining 2% of designs. Thus, each design was assigned a single predicted stability score but appeared as a training sample in 49 other random forest models.

Next, we identified outlier designs that the model predicted to have a high stability score, but actually had a low experimentally measured stability score. Specifically, we ranked designs based on the signed difference in their predicted vs. experimental stability score (= predicted - experimental), and selected the 50 designs with the most positive signed difference. Among this pool, we selected designs with trypsin and chymotrypsin $EC_{50}$ values less than 1 in replicate 1 and 1.5 in replicate 2, an experimental stability score less than 0.3, and a predicted stability score of more than 1. These criteria produced a list of the 21 outlier designs (14 HHH, 4 EHEE, and 3 EEHEE) that we characterized using DMS. See https://github.com/Haddox/design_guided_optE/tree/main/id_outlier_designs for model predictions and experimental stability scores for all designs, as well as a list of all 21 outlier designs characterized by DMS.

S8 Fig illustrates at a mechanistic level how a random forest model works and how its underlying structure helped identify outlier designs. Each random forest model comprised a large collection of decision-trees trained on a point cloud of designs in *n*-dimensional space, where each point is a design and each dimension is a biophysical feature. Each decision tree begins by choosing a single feature, *x*, from a randomly selected subset of the features, along which to bisect the cloud into two subsets. The value of the *x* coordinate at which the cloud is split (call it $x_s$) is chosen to optimally reduce a measure of disparity between the stability scores of all the designs in each half of the cloud, i.e., those designs having feature $x$ values less than $x_s$ and those having $x$ values greater than $x_s$. The bisection of feature space into two regions along the $x$ dimension corresponds to a splitting of the root node in a tree into two branches, and likewise a splitting of the data into two subsets–one associated to each branch. Each subset of designs may again be divided along some feature coordinate at an optimally chosen value, thus dividing feature space into a total of $k \leq 4$ regions, and resulting in a depth-2 tree with $k$ leaves. The process is continued until a stopping criteria is achieved. The result is a binary tree whose terminal leaves encode for disjoint regions of the feature space and the subsets of designs that occupy each region.

The model-predicted stability score for a design, *P*, is taken to be the average stability score of all the other designs appearing in the same regions of feature space (leaf nodes) to which *P* belongs, across all decision trees in the forest. In this way, the random forest models learn the discriminating features and regions of feature space that align with the experimental stability scores, and determine a model-driven notion of proximity between designs that can be used to identify designs which are observed to be unstable but whose neighbors (members of common leaves in the random forest decision trees) are highly stable. Figure S8 shows an example of an outlier design which is unstable but is in the same leaf node of the decision tree as many highly stable designs.

### References


1. Chen Z, Boyken SE, Jia M, Busch F, Flores-Solis D, Bick MJ, et al. Programmable design of orthogonal protein heterodimers. Nature. 2019;565: 106–111.

2. Hicks DR, Kennedy MA, Thompson KA, DeWitt M, Coventry B, Kang A, et al. De novo design of protein homodimers containing tunable symmetric protein pockets. Proceedings of the National Academy of Sciences. 2022;119: e2113400119.

3. Brunette TJ, Parmeggiani F, Huang P-S, Bhabha G, Ekiert DC, Tsutakawa SE, et al. Exploring the repeat protein universe through computational protein design. Nature. 2015;528: 580–584.

4. Xu C, Lu P, Gamal El-Din TM, Pei XY, Johnson MC, Uyeda A, et al. Computational design of transmembrane pores. Nature. 2020;585: 129–134.

5. Boyken SE, Chen Z, Groves B, Langan RA, Oberdorfer G, Ford A, et al. De novo design of protein homo-oligomers with modular hydrogen-bond network–mediated specificity. Science. 2016 [cited 2 May 2025]. doi:10.1126/science.aad8865

6. Lajoie MJ, Boyken SE, Salter AI, Bruffey J, Rajan A, Langan RA, et al. Designed protein logic to target cells with precise combinations of surface antigens. Science. 2020 [cited 2 May 2025]. doi:10.1126/science.aba6527

7. Huddy TF, Hsia Y, Kibler RD, Xu J, Bethel N, Nagarajan D, et al. Blueprinting extendable nanomaterials with standardized protein blocks. Nature. 2024;627: 898–904.

8. Wu K, Bai H, Chang Y-T, Redler R, McNally KE, Sheffler W, et al. De novo design of modular peptide-binding proteins by superhelical matching. Nature. 2023;616: 581–589.

9. Sahtoe DD, Praetorius F, Courbet A, Hsia Y, Wicky BIM, Edman NI, et al. Reconfigurable asymmetric protein assemblies through implicit negative design. Science. 2022 [cited 2 May 2025]. doi:10.1126/science.abj7662

10. Basanta B, Bick MJ, Bera AK, Norn C, Chow CM, Carter LP, et al. An enumerative algorithm for de novo design of proteins with diverse pocket structures. Proceedings of the National Academy of Sciences. 2020;117: 22135–22145.

11. Dou J, Vorobieva AA, Sheffler W, Doyle LA, Park H, Bick MJ, et al. De novo design of a fluorescence-activating β-barrel. Nature. 2018;561: 485–491.

12. Park H, Bradley P, Greisen P Jr, Liu Y, Mulligan VK, Kim DE, et al. Simultaneous Optimization of Biomolecular Energy Functions on Features from Small Molecules and Macromolecules. J Chem Theory Comput. 2016;12: 6201–6212.

13. Pedregosa F, Varoquaux G, Gramfort A, Michel V, Thirion B, Grisel O, et al. Scikit-learn: Machine Learning in Python. Journal of Machine Learning Research. 2011;12: 2825–2830.

14. Rocklin GJ, Chidyausiku TM, Goreshnik I, Ford A, Houliston S, Lemak A, et al. Global analysis of protein folding using massively parallel design, synthesis, and testing. Science. 2017;357: 168–175.

15. Singer JM, Novotney S, Strickland D, Haddox HK, Leiby N, Rocklin GJ, et al. Large-scale

design and refinement of stable proteins using sequence-only models. PLoS One. 2022;17: e0265020.
